# Apical transport of Crumbs maintains epithelial cell polarity

**DOI:** 10.1101/592311

**Authors:** M Aguilar-Aragon, G Fletcher, BJ Thompson

## Abstract

Crumbs (Crb in *Drosophila*; CRB1-3 in mammals) is a transmembrane determinant of epithelial cell polarity and a regulator of Hippo signalling. Crb is normally localized to apical cell-cell contacts, just above adherens junctions, but how apical trafficking of Crb is regulated in epithelial cells remains unclear. We use the *Drosophila* follicular epithelium to demonstrate that polarized trafficking of Crb is mediated by transport along microtubules by the motor protein Dynein and along actin filaments by the motor protein Myosin-V (MyoV). Blocking transport of Crb-containing vesicles by Dynein or MyoV leads to accumulation of Crb within Rab11 endosomes, rather than apical delivery. The final steps of Crb delivery and stabilisation at the plasma membrane requires the exocyst complex and three apical FERM domain proteins – Merlin, Moesin and Expanded – whose simultaneous loss disrupts apical localization of Crb. Accordingly, a knock-in deletion of the Crb FERM-binding motif (FBM) also impairs apical localization. Finally, overexpression of Crb challenges this system, creating a sensitized background to identify components involved in cytoskeletal polarization, apical membrane trafficking and stabilisation of Crb at the apical domain.

## Introduction

Cell polarity is a fundamental characteristic of living organisms.The molecular determinants of cell polarity have been revealed through pioneering genetic screens in the yeasts *S. cerevisiae* and *S. pombe*, the worm *C. elegans*, and the fruit fly *Drosophila melanogaster* (St Johnston and Ahringer, 2010; Tepass, 2012; Thompson, 2013).In yeast, the small GTPase Cdc42 was discovered to be a fundamental determinant of cell polarity (Adams et al., 1990), localizing to one pole of the cell through a positive feedback loop of self-recruitment (Johnson et al., 2011; Martin, 2015; Slaughter et al., 2009). Two general mechanisms for Cdc42-driven positive feedback were identified in yeast: (1) oligomeric clustering of Cdc42 complexes (Altschuler et al., 2008; Bendezu et al., 2015; Irazoqui et al., 2003) and (2) actin cytoskeleton mediated delivery of Cdc42 containing vesicles by a Myosin motor protein (Lechler et al., 2000; Wedlich-Soldner et al., 2003) in *S. cerevisiae* or microtubule mediated transport of polarizing factors in *S. pombe* (Martin and Arkowitz, 2014; Martin et al., 2005; Mata and Nurse, 1997; Minc et al., 2009). In fertilized worm oocytes (zygotes), polarization depends on clustering of Cdc42 and PAR-3 complexes (Dickinson et al., 2017; Gotta et al., 2001; Rodriguez et al., 2017; Sailer et al., 2015). The actin cytoskeleton flows to one pole of the worm zygote, pulling Cdc42-containing PAR-3 complexes along with it via bulk fluid ‘advection’ (Goehring et al., 2011). However, the actin cytoskeleton is not strictly required for polarization of the worm zygote, which can also be triggered by a microtubule-based mechanism (Motegi et al., 2011; Zonies et al., 2010). In *Drosophila* and mammalian oocytes, the actin cytoskeleton is polarized and plays an important role with Cdc42 in breaking symmetry (Leblanc et al., 2011; Leibfried et al., 2013; Ma et al., 2006; Wang et al., 2013; Yi et al., 2013; Yi et al., 2011; Zhang et al., 2008). In epithelial cells, which of these two mechanisms, oligomeric clustering of determinants versus cytoskeletal transport of determinants, is responsible for directing polarity remains a fundamental unsolved problem.

*Drosophila* epithelial cells exhibit a more complex polarization than oocytes, with the plasma membrane divided into distinct apical and basolateral domains, separated by a ring of adherens junctions (Gibson and Perrimon, 2003; St Johnston and Ahringer, 2010; Tepass, 2012; Thompson, 2013). Like oocytes, epithelial cells express the cortical polarity determinant Par-3/Bazooka (Baz), which is polarized through oligomeric clustering at the plasma membrane (Benton and St Johnston, 2003a, b; Harris, 2017; Krahn et al., 2010; McKinley et al., 2012; Mizuno et al., 2003). In addition, epithelial cells express a second transmembrane polarity determinant Crumbs (Crb) (Bazellieres et al., 2018; Campbell et al., 2009; Grawe et al., 1996; Knust et al., 1993; Tepass, 2012; Tepass and Knust, 1993; Tepass et al., 1990). Baz and Crb act in parallel to maintain the apical domain via recruitment of the Cdc42-Par6-aPKC complex (Fletcher et al., 2015; Fletcher et al., 2012; Hutterer et al., 2004; Joberty et al., 2000; Petronczki and Knoblich, 2001; Shahab et al., 2015; Tanentzapf and Tepass, 2003). The Cdc42-Par6-aPKC complex promotes Crb polarization, forming a positive feedback loop whose nature is still not fully understood (Fletcher et al., 2012; Harris and Tepass, 2008). The redundancy between Baz and Crb makes it possible to study Crb polarization in isolation, because defects in Crb localization to the apical domain do not disrupt the overall apical-basal polarization of most epithelial cells, and instead disrupts Hippo signalling (Chen et al., 2010; Fletcher et al., 2015; Fletcher et al., 2012; Ling et al., 2010) – indeed the crucial requirement for Crb in epithelial polarity occurs during embryonic gastrulation (Campbell et al., 2009; Grawe et al., 1996), when Baz is localized to adherens junctions in a planar polarized fashion during germ-band extension (Simoes Sde et al., 2010; Zallen and Wieschaus, 2004).

We previously proposed a model of Crb localization via Cdc42-dependent positive feedback, based on analysis of overexpressed Crb in *Drosophila* follicle cells (Fletcher et al., 2012). In this model, Crb is delivered apically from Rab11 endosomes (Blankenship et al., 2007; Li et al., 2007; Roeth et al., 2009) and engages in oligomeric clustering via its extracellular domain, as well as interacting via its cytoplasmic domain with the PDZ domain protein Stardust/PALS1 [Sdt (Bachmann et al., 2001; Hong et al., 2001; Knust et al., 1993; Muller and Wieschaus, 1996; Tepass and Knust, 1993)] and the FERM domain proteins Moesin [Moe (Medina et al., 2002; Polesello et al., 2002; Sherrard and Fehon, 2015b; Speck et al., 2003)] and Expanded [Ex (Chen et al., 2010; Ling et al., 2010)] to regulate Crb localization at the plasma membrane (Fletcher et al., 2012; Thompson et al., 2013). It was further suggested that aPKC phosphorylates the Crb FERM-binding domain (Sotillos et al., 2004) and that this domain might contribute to stabilization of Crb at the apical domain (Fletcher et al., 2012). Further genetic and structural evidence supports a key role for direct interaction with Sdt/PALS1 in maintaining endogenous Crb at the plasma membrane by preventing Crb endocytosis (Ivanova et al., 2015; Li et al., 2014; Lin et al., 2015) and that aPKC phosphorylation of Crb may antagonize Moe binding in favour of Ex (Sherrard and Fehon, 2015b; Su et al., 2017) or Sdt (Wei et al., 2015). However, recent evidence indicates that deletion of the extracellular domain of endogenously expressed Crb, or mutation of the FERM binding domain of endogenously expressed Crb, does not disrupt epithelial polarity during embryonic gastrulation in *Drosophila* (Cao et al., 2017; Das and Knust, 2018; Klose et al., 2013), although it can affect Crb localization in the follicle cell epithelium (Sherrard and Fehon, 2015a).Furthermore, FERM domain phosphorylation is also not strictly essential for endogenous Crb localization or function in the embryo (Cao et al., 2017). The reasons for the discrepancy in results between overexpressed Crb (Fletcher et al., 2012; Letizia et al., 2013; Letizia et al., 2011; Roper, 2012) and endogenous Crb (Cao et al., 2017; Das and Knust, 2018) remain unclear and are important to resolve. Evidently, mechanisms of positive feedback exist that can recruit Crb to the apical domain in the absence of Crb-Crb oligomeric clustering via the extracellular domain or aPKC phosphorylation of the Crb intracellular domain.

We therefore considered alternative mechanisms of positive feedback that might promote Crb localization to the apical domain. Since Crb is a transmembrane protein, it is conceivable that its exocytic delivery to the plasma membrane could become polarized by directional transport along either microtubules or actin.Microtubule polarity depends on core epithelial polarity determinants, which localize the microtubule minus-end binding proteins Shot (MACF1, ACF7, BPAG1 in mammals) and Patronin (CAMSAP1, CAMSAP2, CAMSAP3 in mammals) for acentrosomal nucleation apically (Khanal et al., 2016; Noordstra et al., 2016; Toya et al., 2016). The actin cytoskeleton is organized by the Rho GTPase and adherens junctions (Cox et al., 1996; Fox et al., 2005; Homem and Peifer, 2008; Muller and Wieschaus, 1996; Peifer et al., 1993) which are ultimately positioned between the core apical and basolateral polarity determinants. The apical determinants include Baz (Muller and Wieschaus, 1996), Crb (Grawe et al., 1996; Tanentzapf et al., 2000; Tepass et al., 1990), Sdt (Bachmann et al., 2001; Hong et al., 2001; Knust et al., 1993; Muller and Wieschaus, 1996; Tepass and Knust, 1993), Cdc42 (Atwood et al., 2007; Eaton et al., 1995; Genova et al., 2000; Harris and Tepass, 2008; Hutterer et al., 2004; Joberty et al., 2000; Peterson et al., 2004) and its effector kinases aPKC (Hutterer et al., 2004; Kim et al., 2009; Wodarz et al., 2000), Pak1 (Aguilar-Aragon et al., 2018; Conder et al., 2007; Harden et al., 1996), Pak4 (Walther et al., 2016), Gek/MRCK (Luo et al., 1997; Zihni et al., 2017) as well as WASP (Leibfried et al., 2008) and the Arp2/3 complex (Georgiou et al., 2008; Leibfried et al., 2013).The basolateral polarity determinants Lgl, Scrib, Dlg and Yurt engage in mutual antagonism with apical polarity determinants and are thus necessary for cytoskeletal polarization (Bilder et al., 2000; Bilder and Perrimon, 2000; Bilder et al., 2003; Gamblin et al., 2014; Laprise et al., 2006; Laprise et al., 2009; Tanentzapf et al., 2000; Tanentzapf and Tepass, 2003).Despite this wealth of understanding of how the cytoskeleton is polarized along the apical-basal axis in epithelia, whether it might function to direct membrane trafficking of Crb remains poorly understood.

Here we show that the actin motor protein Myosin-V (MyoV), previously implicated in Rab11-mediated apical secretion and found to co-localize with Crb (Li et al., 2007; Pocha et al., 2011), as well as the microtubule motor protein Dynein, previously implicated in transporting *crb* and *sdt* mRNA (Cao et al., 2017; Horne-Badovinac and Bilder, 2008; Li et al., 2008), both function in directing membrane trafficking of the Crb protein from Rab11 endosomes (Fletcher et al., 2012; Roeth et al., 2009) to the apical domain of the ovarian follicle cell epithelium. Accordingly, disruption of microtubule polarization and/or the actin cytoskeleton impairs Crb localization to the apical domain. We confirm that the Exocyst complex then promotes the final step in delivery of Crb to the plasma membrane, as previously demonstrated in the embryo (Blankenship et al., 2007). Crb must then interact with apical FERM-domain proteins via its FERM-binding motif to be stabilized at the plasma membrane. Finally, we find that overexpression of Crb challenges this system of polarized transport and exocytic delivery, acting as a sensitized background for the testing of molecular mechanisms involved in Crb polarization.

## Results

We began by re-examining the requirement for the microtubule motor protein Dynein in regulation of Crb in the *Drosophila* ovarian follicle cell epithelium. Previously, loss of Dynein was reported to cause decreased levels of Crb protein in follicle cells due to failure of *crb* or *sdt* mRNA transport (Cao et al., 2017; Horne-Badovinac and Bilder, 2008; Li et al., 2008). Using a Crb-GFP knockin line, we find that, in addition to reduced Crb protein at the apical membrane domain, Crb also accumulates basally in Rab11 endosomes upon silencing of Dynein expression by conditional expression of *UAS.dynein-RNAi* hairpins with the follicle cell-specific *trafficjam.Gal4* (*tj.Gal4*) driver (Fig 1A,B; Fig 1-S1). We find similar results with antibody staining for Crb, which localizes basally upon clonal expression of *actin>flipout>Gal4 UAS.dynein-RNAi* hairpins but not in neighbouring wild-type follicle cells (Fig 1C). Co-staining for Crb and Rab11 in wild-type and *dynein-RNAi* expressing follicle cells reveals that Crb protein accumulates within Rab11-positive endosomes (Fig 1D). These results suggest that upon loss of the microtubule minus-end directed motor protein Dynein, Rab11 endosomes containing Crb protein are transported basally, rather than apically.

**Figure 1.**
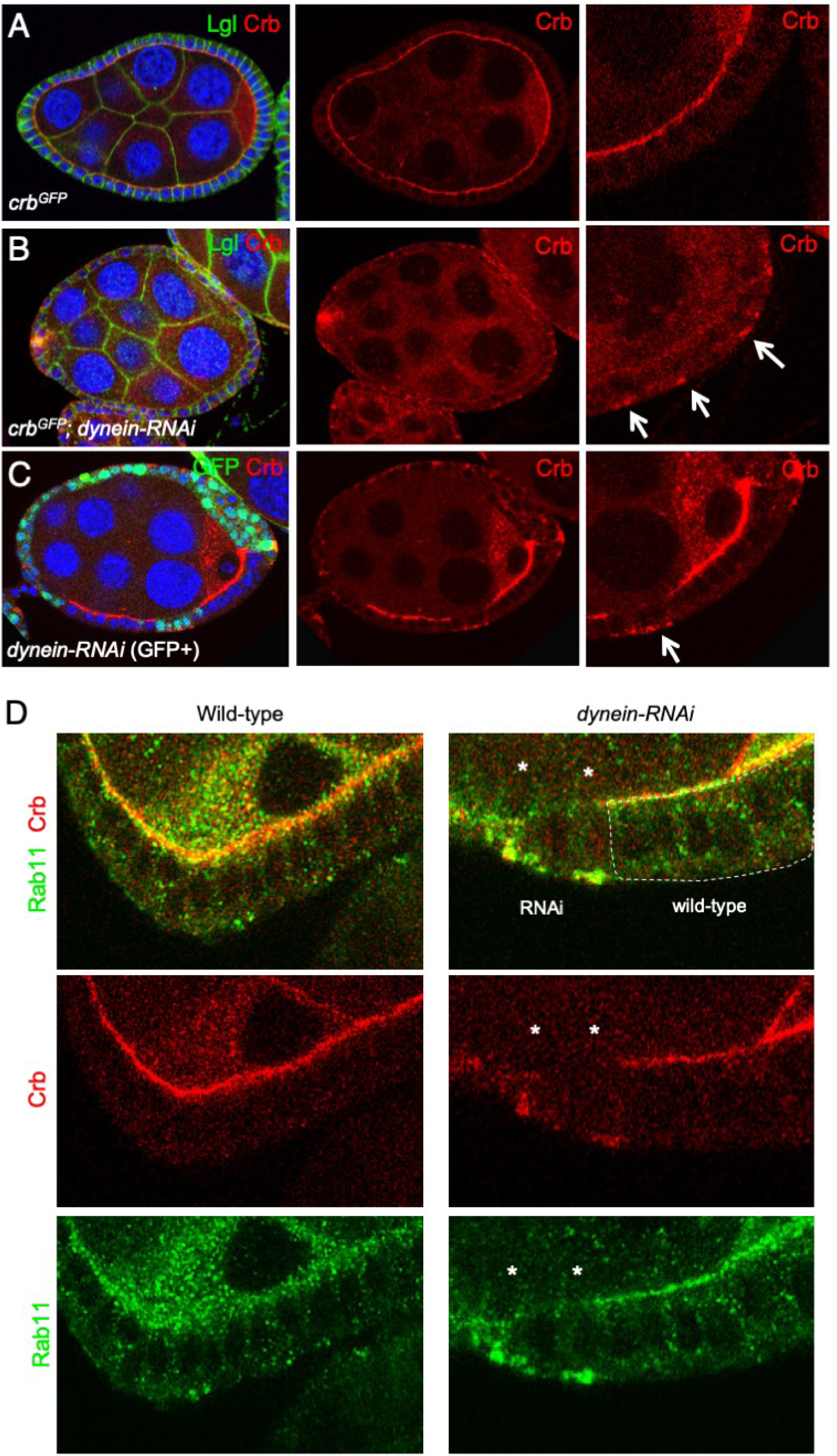
Dynein is required to traffic Crumbs protein apically in the *Drosophila* follicle cell epithelium. A) Endogenously tagged knockin Crb-GFP localises apically in the follicle cell epithelium. B) Expression of Dynein hairpin RNAi throughout the follicle cell epithelium with tj.Gal4 causes relocalisation of Crb-GFP to the basal surface of follicle cells. C) Clonal expression of Dynein hairpin RNAi, marked by expression of nls-GFP, which causes cell-autonomous relocalisation of Crb to the basal surface of follicle cells within the clone D) Co-immunostaining for Crb and Rab11 upon clonal expression of Dynein hairpin RNAi, marked by expression of nls-GFP, reveals co-localization within basal endosomes.

The basal transport of endosomes is normally mediated by the plus-end directed microtubule motor protein Kinesin. We therefore sought to confirm that, in the absence of Dynein, Kinesin is responsible for basal transport of Crb. We find that the basal transport of Crb-GFP in *dynein-RNAi* expressing follicle cells is partially rescued by simultaneous expression of *kinesin-RNAi* (Fig 2A-D). These results confirm that both Dynein and Kinesin are capable of transporting Crb containing Rab11-positive endosomes, but that under normal conditions the Dynein-dependent transport is more efficient, leading to apical localization of Rab11 endosomes and delivery of the Crb protein to the apical plasma membrane domain.

**Figure 2.**
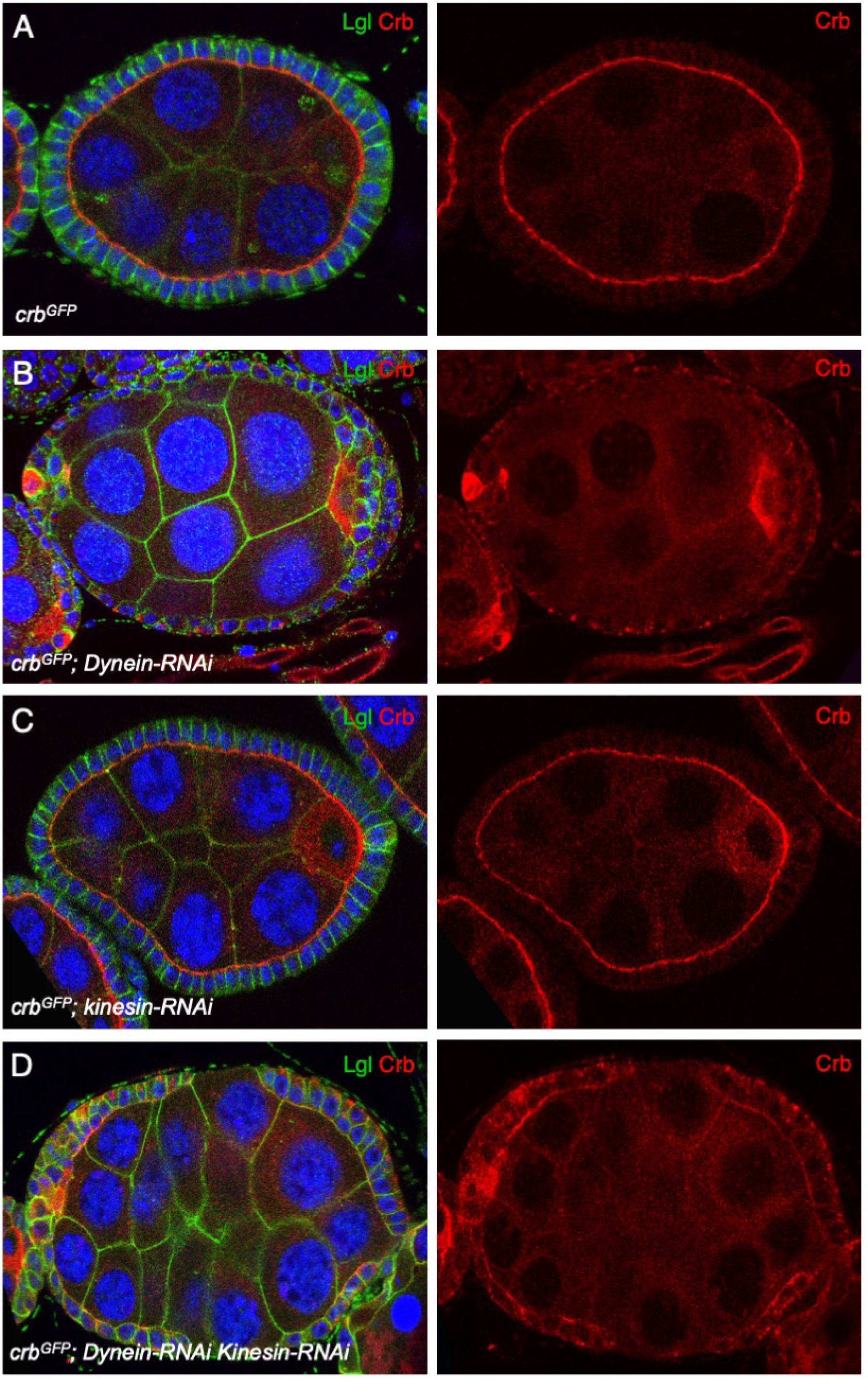
Dynein loss is partially rescued by Kinesin-RNAi. A) Endogenously tagged knockin Crb-GFP localises apically in the follicle cell epithelium. B) Expression of Dynein hairpin RNAi throughout the follicle cell epithelium with tj.Gal4 causes relocalisation of Crb-GFP to the basal surface of follicle cells. C) Expression of Kinesin hairpin RNAi throughout the follicle cell epithelium with tj.Gal4 does not affect localisation of Crb-GFP in follicle cells. D) Expression of both Dynein and Kinesin hairpin RNAi throughout the follicle cell epithelium with tj.Gal4 does partially rescues the effect of Dynein RNAi on localisation of Crb-GFP in follicle cells.

In addition to transport along microtubules by the Dynein motor, it is conceivable that Crb may also be transported along actin filaments by motors such as Myosin V (MyoV), which has been previously shown to form a complex with Crb and to promote apical secretion from Rab11 endosomes in the *Drosophila* photoreceptor (Li et al., 2007; Pocha et al., 2011). Class V Myosins were originally discovered in the yeast *S. cerevisiae* to promote polarized transport along F-actin filaments (Johnston et al., 1991; Pruyne et al., 2004; Pruyne et al., 1998), alongside Class I Myosins (Anderson et al., 1998; Goodson et al., 1996; Lechler et al., 2001; Lechler et al., 2000; Wedlich-Soldner et al., 2003). We find that MyoV-GFP also localizes apically in the follicle cell epithelium, while a dominant-negative form of MyoV named MyoV-GT-GFP (Krauss et al., 2009) fails to localize apically and instead localizes on endosomes (Fig 3A). Expression of MyoV-GT-GFP prevents delivery of Crb to the apical membrane, such that the Crb protein is trapped inside large endosomes that reside just underneath the apical domain (Fig 3B,C). The apical localization of these enlarged Crb-positive endosomes is completely disrupted upon co-expression of *UAS.dynein-RNAi*, which leads to basal localization (Fig 3D,E). Co-staining of Crb and Rab11 confirms that these Crb-containing endosomes are indeed Rab11-positive (Fig 3F,G). These results confirm a dual requirement for transport of the Crb protein by F-actin and microtubule motor proteins.

**Figure 3.**
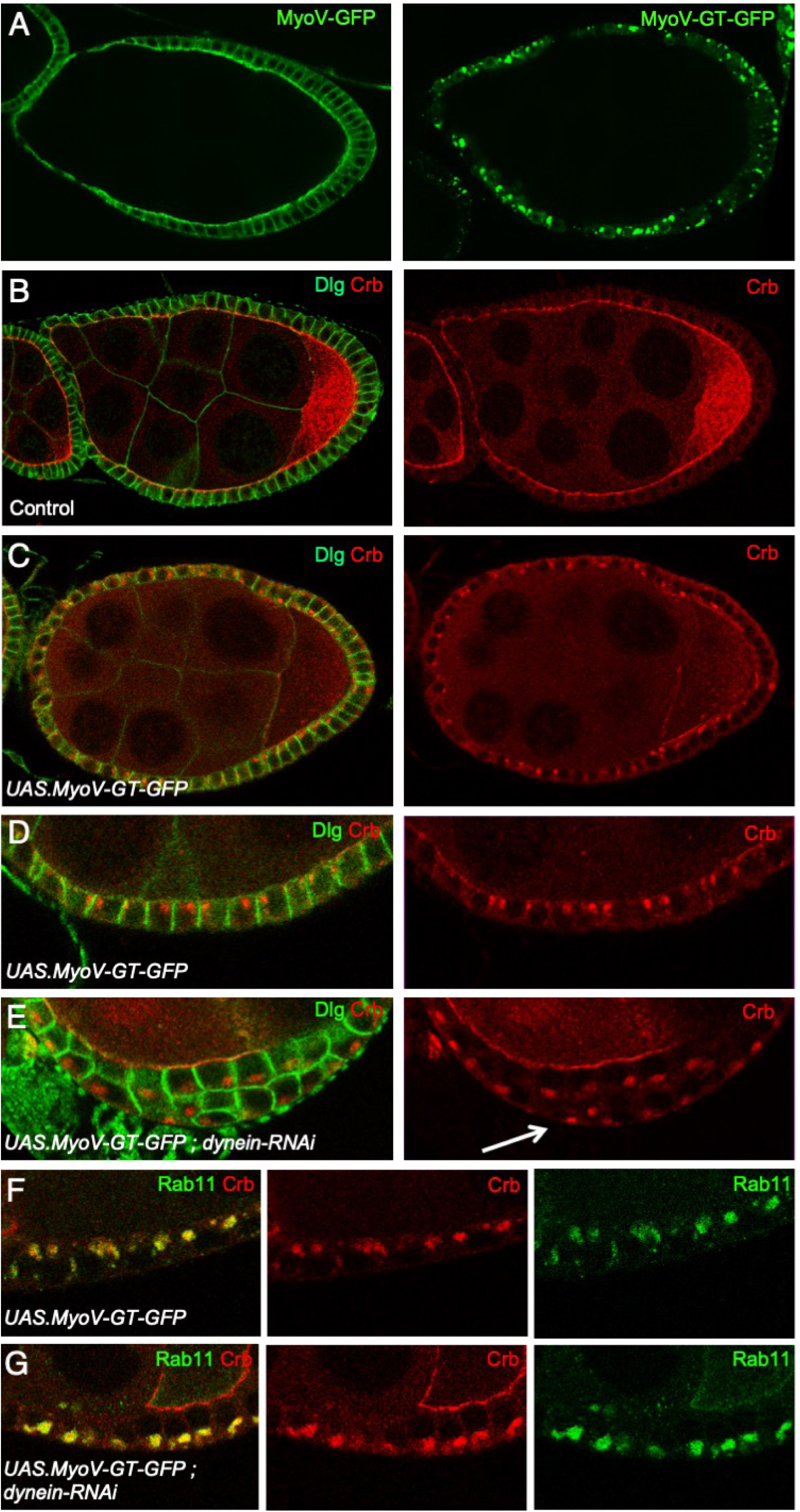
Dominant-negative Myosin V prevents apical delivery of Crumbs, revealing apical transport of entire Crumbs-containing Rab11-positive endosomes by Dynein. A) GFP-tagged MyoV expressed with tj.Gal4 localises primarily to the apical domain of follicle cells, while GFP-tagged MyoV GT (dominant-negative) localises to endosomes (n>10 egg chambers). B) Control egg chambers immunostained for Dlg and Crb (n>12 egg chambers). C) Expression of dominant-negative MyoV GT prevents apical localisation of Crb, which instead accumulates in endosomes that localise towards the apical pole of the cell (n>15 egg chambers). D) High magnification view of C (n>15 egg chambers). E) Co-expression of Dynein hairpin RNAi with dominant-negative MyoV GT causes accumulation of Crb in endosomes that localise basally (n>11 egg chambers). F) Co-immunostaining for Rab11 and Crb upon expression of dominant-negative MyoV GT reveals co-localisation in enlarged apical endosomes (n>8 egg chambers). G) Co-immunostaining for Rab11 and Crb upon expression of dominant-negative MyoV GT and dynein-RNAi hairpins reveals co-localisation in enlarged basal endosomes (n>9 egg chambers).

Given our findings with MyoV, we sought to confirm that the F-actin cytoskeleton is also necessary for apical localization of Crb. As expected, disruption of F-actin by acute treatment with Latrunculin A caused a strong loss of MyoV-GFP and Crb-GFP from the apical domain, with Crb-GFP accumulating in endosomes (Fig 4A-F). Treatment with another F-actin cytoskeleton disrupting compound, Cytochalasin D, had a similar effect on Crb-GFP localization (Fig 3G). These results confirm the essential requirement for the F-actin cytoskeleton in polarization of Crb.

**Figure 4.**
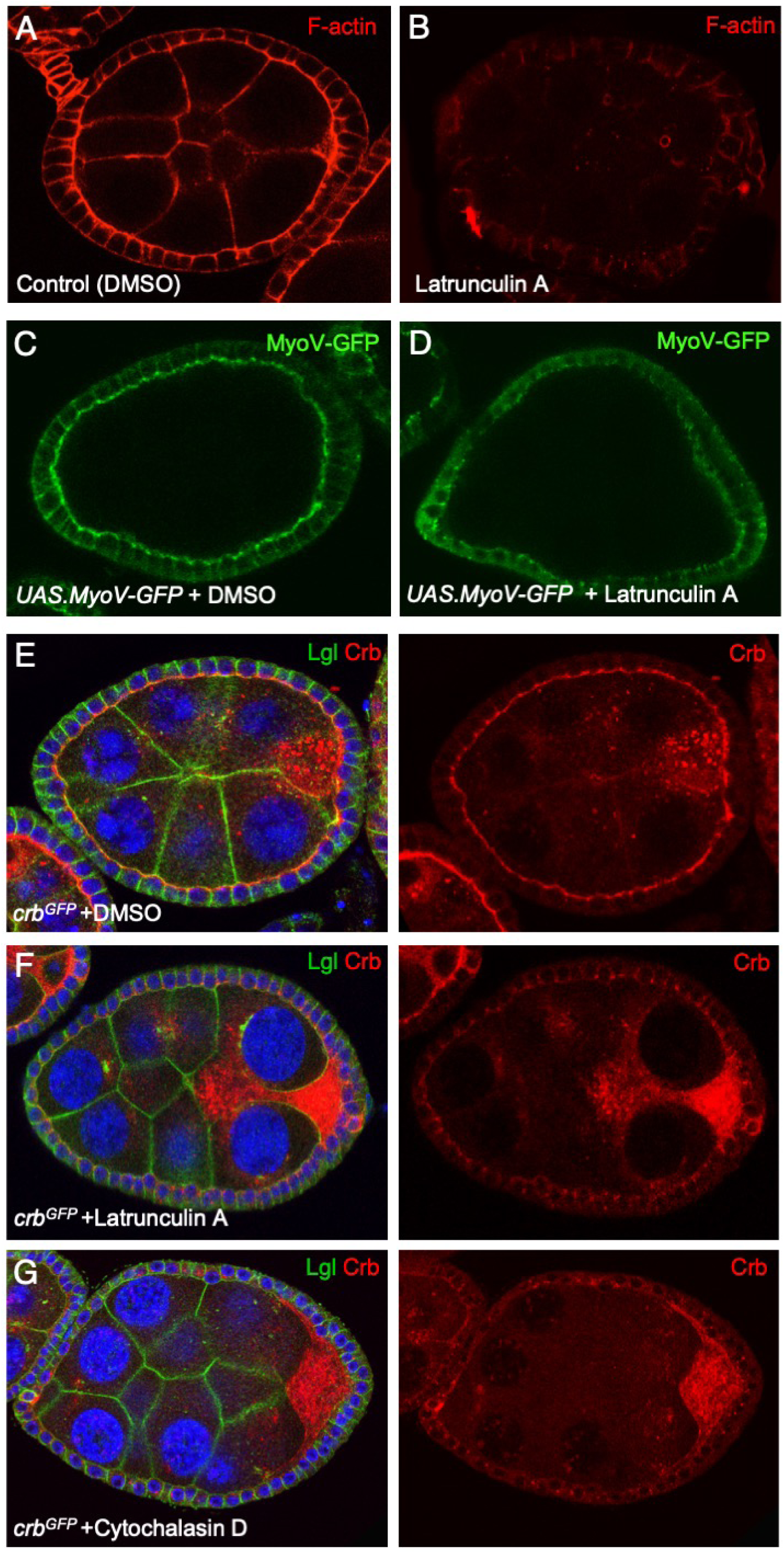
The F-actin cytoskeleton, which concentrates apically, is required for apical localisation of both MyoV and Crb. A) Control (DMSO-treated) egg chamber stained for F-actin with phalloidin (n>9 egg chambers). B) Latrunculin A treated egg chamber stained for F-actin with phalloidin (n>10 egg chambers). C) Control (DMSO-treated) egg chamber expressing MyoV-GFP with tj.Gal4 (n>10 egg chambers). D) Latrunculin A treated egg chamber expressing MyoV-GFP with tj.Gal4 shows loss of MyoV-GFP apical localisation (n>12 egg chambers). E) Control (DMSO-treated) Crb-GFP egg chamber showing normal apical localisation of Crb in follicle cells (n>7 egg chambers). F) Latrunculin A treated Crb-GFP egg chamber showing loss of apical Crb localisation and localisation to endosomal punctae (n>5 egg chambers). G) Cytochalasin D treated Crb-GFP egg chamber showing loss of apical Crb localisation and localisation to endosomal punctae (n>7 egg chambers). H) Jasplakinolide treated Crb-GFP egg chamber showing loss of apical Crb localisation and localisation to endosomal punctae (n>12 egg chambers).

We next sought to examine the role of the apically-localized FERM domain proteins – Merlin (Mer), Expanded (Ex) and Moesin (Moe). FERM domains link the actin cytoskeleton to the plasma membrane (Chishti et al., 1998) and bind directly to spectrins (Baines et al., 2014), which are required to polarize microtubules and regulate the Hippo pathway in *Drosophila* (Fletcher et al., 2015; Khanal et al., 2016). Mer and Ex are redundantly required to regulate the Hippo signaling pathway in *Drosophila*, a parallel function that may arise from the different subcellular localizations of Mer, which is found across the apical surface, and Ex, which localizes to the sub-apical junction through direct interaction with the Crb intracellular domain (Fletcher et al., 2015; Hamaratoglu et al., 2006; Su et al., 2017). Moe has important roles in linking cortical F-actin to the plasma membrane, particularly during mitotic cell rounding and microvilli formation (Carreno et al., 2008; Fehon et al., 2010; Kunda et al., 2008; Sauvanet et al., 2015). We find that mutation of *moe* does not affect Crb localization in follicle cells, similar to *mer, ex* double mutants (Fig 5A,B). Double mutants of *mer, moe* also have no effect on Crb (Fig 5C). However, triple mutant clones of *mer, ex, moe* do lead to a strong disruption of Crb localization (Fig 5D). Accordingly, knock-in deletion of the Crb FERM-binding motif (FBM) also reduces the apical localization of Crb in follicle cells (Fig 5E,F). These findings demonstrate that the apical FERM domain proteins are collectively required to promote Crb polarization, a role that is consistent with their molecular functions in polarising the cytoskeleton in epithelia and with their ability to directly bind the Crb intracellular domain.

**Figure 4.**
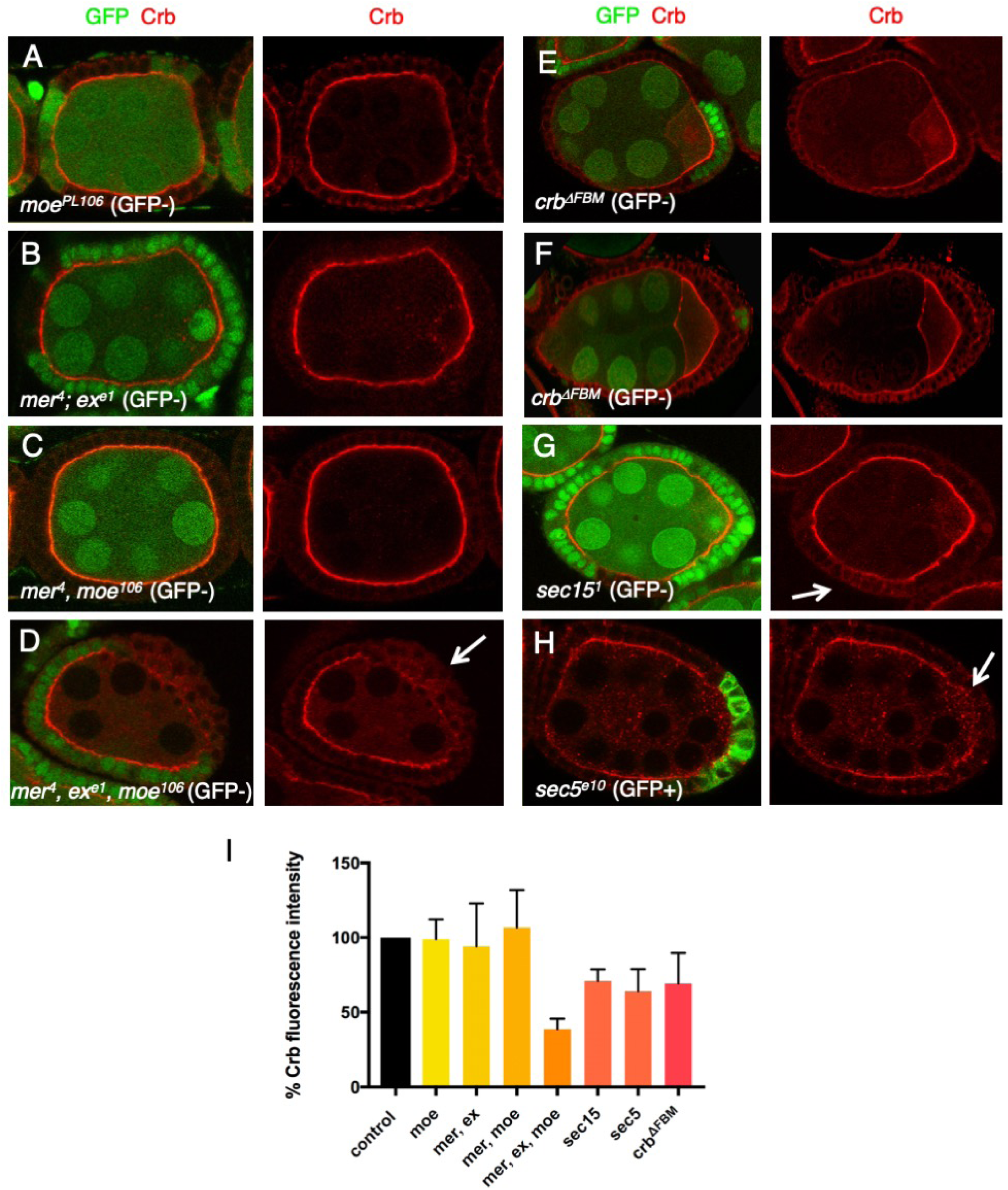
Apical FERM domain proteins and the Exocyst promote apical delivery of Crb. A) Mutant clones for *moesin*, marked by absence of GFP, show normal Crb localisation (n>12 egg chambers). B) Double mutant clones for *merlin* and *expanded*, marked by absence of GFP, show normal Crb localisation (n>6 egg chambers). C) Double mutant clones for *merlin* and *moesin*, marked by absence of GFP, show normal Crb localisation (n>14 egg chambers). D) Triple mutant clones for *merlin, expanded* and *moesin*, marked by absence of GFP, show loss of apical Crb localisation (n>4 egg chambers). E) Mutant clones for *sec15*, marked by absence of GFP, show loss of apical Crb localisation (n>5 egg chambers). F) Mutant clones for *sec15*, marked positively by expression of GFP, show loss of apical Crb localisation in extruded cells (n>7 egg chambers).

Once Crb has been successfully trafficked to the apical membrane of the cell, it must be delivered to the plasma membrane through a process of regulated exocytosis. The exocyst complex was discovered to mediate polarized exocytosis in the yeast *S. cerevisiae* (He and Guo, 2009; Novick et al., 1980; TerBush et al., 1996). The exocyst subunit Sec15 directly interacts with the Cdc42-mediated polarity establishment complex in yeast (France et al., 2006). In mammalian epithelial cells in culture, the exocyst associates with adherens junctions, tight junctions (Yeaman et al., 2004), and Par3 (Ahmed and Macara, 2017). In *Drosophila* epithelia, the exocyst component Exo84 is required for apical trafficking of Crb from Rab11 endosomes to the plasma membrane in embryos (Blankenship et al., 2007); Sec6 is required for apical exocytosis in *Drosophila* photoreceptors (Beronja et al., 2005); and Sec5 is required for efficient delivery of E-cadherin to adherens junctions in the pupal notum (Langevin et al., 2005) but is not required for cytoskeletal polarization in follicle cell epithelium (Murthy and Schwarz, 2004). We therefore tested whether Sec15 and Sec5 were required for apical localization of Crb in the follicle cell epithelium. We find that mutants clones for *sec15* or *sec5* strongly disrupt the apical localization of Crb (Fig 4G-I). These findings confirm an essential requirement for the exocyst in delivery of Crb to the apical membrane in the follicular epithelium.

Finally, we sought to challenge the system of polarized Crb exocytosis by overexpressing full-length Crb protein in the follicle cell epithelium. We find that this results in ectopic localization of Crb to lateral membranes, in addition to the apical domain (Fig 6A,B). Importantly, loss of individual components of the cytoskeletal polarization/trafficking machinery or exocyst dramatically enhances the Crb overexpression phenotype, leading to massive accumulation of Crb in endosomes (Fig 6A,B). This genetic interaction also occurs for genes whose individual loss-of-function has no detectable effect on Crb in a wild-type background, such as *patronin* or *shot*. The assay further highlights the key role of adherens junctions, whose disruption with alpha-catenin-RNAi strongly affects the F-actin cytoskeleton and causes a striking endosomal accumulation of overexpressed Crb (Fig 6B). The assay similarly reveals a key role for the Crb extracellular domain, whose loss leads to strong accumulation of the overexpressed Crb transmembrane and intracellular domain (CrbICD) in endosomes, even though endogenously-expressed CrbICD exhibits only a moderate reduction in apical localization (Fig 6C-E). Thus, Crb overexpression in follicle cells is a highly sensitized assay for detecting components of the Crb trafficking and polarization machinery, even where such components exhibit redundancy or degeneracy with others or have pleiotropic phenotypes. A summary of the key components of cytoskeletal polarization and Crb trafficking is shown schematically in the final diagram (Fig 6F).

**Figure 6.**
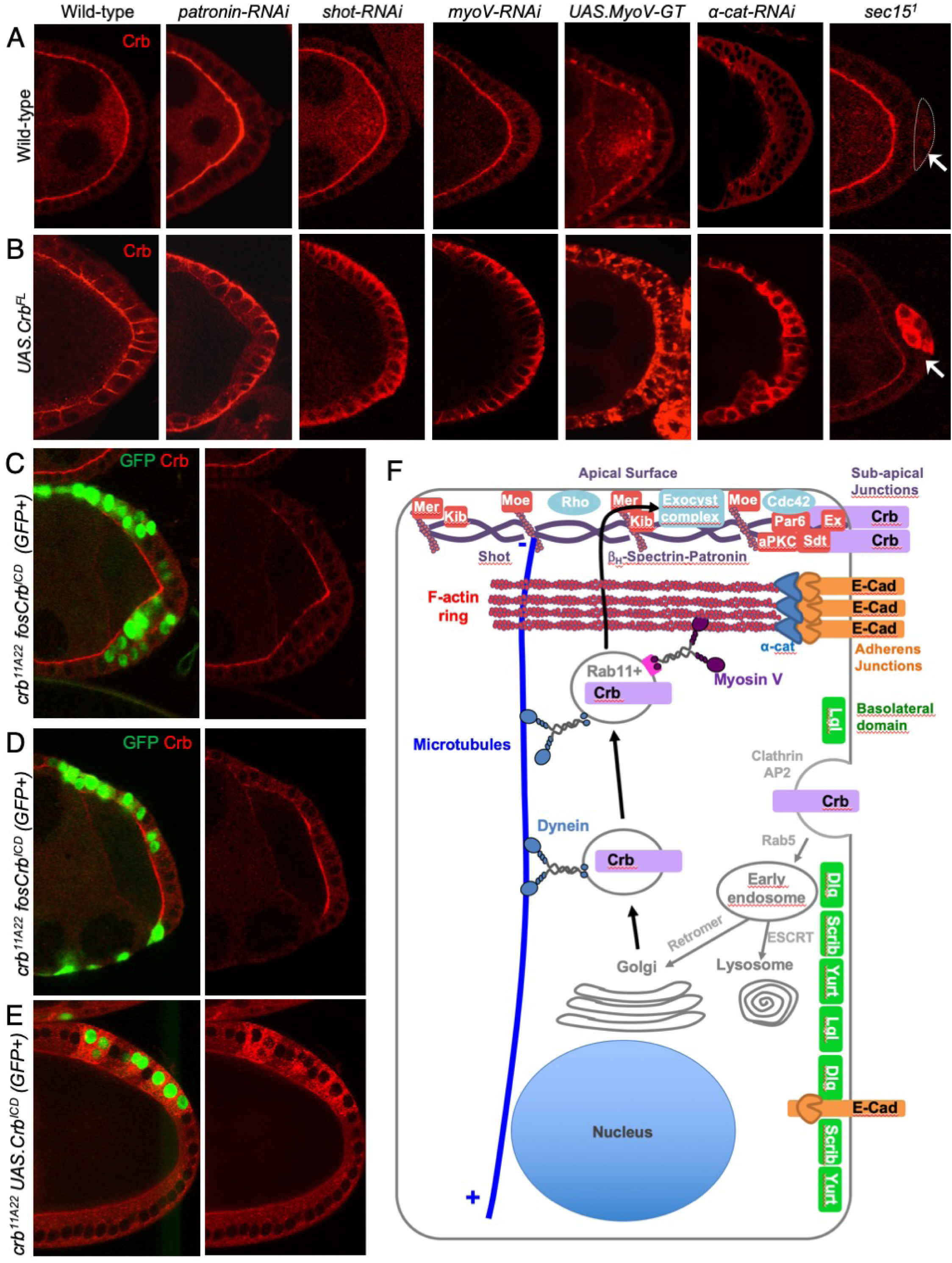
Overexpression of Crumbs is a sensitized assay to identify genes required for its apical trafficking and delivery. A) Crb localisation in wild-type posterior follicle cells and those of the indicated genotypes, either transgenes expressed with tj.gal4 or mutant clones of *sec15* (arrow) (n>8 samples per experimental condition). B) Overexpressed full-length Crb localisation in otherwise wild-type posterior follicle cells and those of the indicated genotypes, either transgenes expressed with tj.gal4 or mutant clones of *sec15* (arrow). Note the strong accumulation of overexpressed Crb upon manipulation of key trafficking regulators, including those that disrupt the microtubule cytoskeleton. Thus, although some factors are normally dispensable for endogenous Crb localisation, they are essential to localise overexpressed Crb. (n>8 samples per experimental condition). C) Null mutant clones for *crb* expressing Crb intracellular domain (ICD) at endogenous levels from a fosmid (marked by expression of GFP). D) Null mutant clones for *crb* expressing Crb intracellular domain (ICD) at endogenous levels from a fosmid (marked by expression of GFP). E) Null mutant MARCM clones for *crb* expressing Crb intracellular domain (ICD) at high levels with the GAL4/UAS system (marked by expression of GFP). F) Schematic diagram of key components of the cellular cytoskeleton and trafficking machinery and their functions in localising Crb to the apical domain of epithelial cells.

## Discussion

Polarization of the cytoskeleton is a universal feature of columnar epithelial cells that determines polarized membrane trafficking of many cargo proteins and is known to depend on fundamental determinants of epithelial polarity (Mellman and Nelson, 2008; Mostov et al., 2000; Nelson, 1991, 2003; Rodriguez-Boulan and Macara, 2014; Weisz and Rodriguez-Boulan, 2009). Whether the localization of apical-basal polarity determinants themselves requires cytoskeletal polarization has been uncertain, particularly because apical-basal polarization of the Baz/Par-3 system can arise without cytoskeletal polarization or membrane trafficking in asymmetrically dividing *Drosophila* neuroblasts or *C. elegans* zygotes (Halbsgut et al., 2011; St Johnston and Ahringer, 2010; Tepass, 2012; Thompson, 2013). Furthermore, disruption of microtubules or preventing polarization of microtubules does not strongly interfere with polarization of the plasma membrane in *S. cerevisiae* (Martin and Arkowitz, 2014), in the *Drosophila* follicle cell epithelium (Khanal et al., 2016) or in cultured mammalian epithelial cells (Noordstra et al., 2016; Toya et al., 2016). In the case of Crb, an apical polarity determinant and transmembrane protein that moves through the secretory pathway via Rab11 endosomes and the exocyst to reach the apical domain (Blankenship et al., 2007; Fletcher et al., 2012; Li et al., 2007; Roeth et al., 2009) (Fig S1), it was previously unclear whether polarized cytoskeletal transport was in fact required for apical membrane trafficking and localization of the Crb protein, rather than *crb* or *sdt* mRNA (Cao et al., 2017; Horne-Badovinac and Bilder, 2008; Li et al., 2008).

Our results demonstrate that cytoskeletal polarization and directed motor transport of Crb protein are necessary for its localization to the apical domain of the *Drosophila* follicular epithelium. Disruption of either Dynein-mediated microtubule transport or MyoV-mediated actin transport leads to trapping of Crb in endosomes, such that it is unable to reach the apical domain (Figs 1-3), without affecting overall epithelial polarity (Fig S1). Microtubules are polarized along the apical-basal axis of epithelial cells, with minus ends apical, such that loss of the minus-end directed motor Dynein leads to abnormal basally-directed transport of Crb endosomes (Fig 1). The F-actin cytoskeleton concentrates apically, such that dominant-negative MyoV traps Crb in endosomes that are still transported towards the apical pole of the cell on microtubules, but appear unable to traverse the thick cortical F-actin at the apical surface to reach the plasma membrane (Figs 3&4). Our findings confirm and extend previous work demonstrating that mutation of exocyst complex components also prevents apical delivery of Crb (Fig 5). Thus, membrane trafficking of Crb occurs by directed motor-driven transport along polarized microtubules and F-actin filaments, followed by exocyst-mediated delivery to the plasma membrane, and is crucial for the apical localization of this key polarity determinant.

Our findings also shed light on the mechanisms of cytoskeletal polarization. We have identified a key role for the apical FERM domain proteins, which link the PIP2-rich plasma membrane with F-actin, apical spectrins and the microtubule minus-end binding proteins Shot and Patronin (Chishti et al., 1998; Fehon et al., 2010; Khanal et al., 2016). This apical cortical meshwork (equivalent to the terminal web in mammalian epithelia) is then responsible for polarizing microtubules and thus ensuring directed apical trafficking of Crb by Dynein. Further work is necessary to understand precisely how the three apical FERM domain proteins become localized, and how the apical meshwork is organized, particularly as Moe and Mer are normally found across the apical surface (with Shot) while Ex is found at the sub-apical junctions (with a β_H_-Spectrin-Patronin complex) (Su et al., 2017). Importantly, cytoskeletal polarization ultimately depends on the core apical-basal polarity determinants, including Crb itself, which acts redundantly with Baz to organize epithelial polarity (Tepass, 2012; Thompson et al., 2013). Thus, there is a positive feedback loop between apical polarity determinants and cytoskeletal polarization, which then directs further delivery of Crb to the same location on the plasma membrane to reinforce apical identity and maintain a polarized cytoskeleton.

The above mechanism of cytoskeletal polarization is required for localization of Crb, a transmembrane protein, but is not required for polarization of Baz, a cytoplasmic protein that associates with the plasma membrane. Thus, during asymmetric division of *Drosophila* neuroblasts, which are polarized along the apical-basal axis by Baz, there is no role for either Crb or membrane trafficking (Halbsgut et al., 2011; Hong et al., 2001). Furthermore, during the early establishment of epithelial polarity in the *Drosophila* embryo, polarity is initiated by Baz before Crb becomes expressed (Blankenship et al., 2007; Harris and Peifer, 2004). It is conceivable that the Baz system is able to polarize more rapidly than the Crb system, which may be an advantage in asymmetric cell division and in early establishment of epithelial polarity, but that the Crb system is advantageous in mature epithelial cells, where stable polarization of the cytoskeleton is fundamental to both cellular structure and function. Importantly, redundancy between Baz and Crb-Sdt in recruiting the Cdc42-Par6-aPKC complex enables one system to maintain apical-basal polarity while the other is deployed in a planar polarized fashion during various episodes of morphogenetic change during development (Campbell et al., 2009; Grawe et al., 1996; Thompson et al., 2013).

Our findings also indicate that overexpression of Crb can saturate this system of polarized transport, such that Crb accumulates abnormally along lateral membranes (Fig 6A,B). Overexpressed Crb can also accumulate dramatically within endosomes when individual components of the polarized transport machinery are compromised (Fig 6A,B). The sensitized background caused by Crb overexpression explains the discrepancy between deletion of the conserved extracellular domain in overexpressed Crb, which has dramatic consequences (Fletcher et al., 2012; Letizia et al., 2013; Roper, 2012; Thompson et al., 2013), versus the same experiment in endogenous Crb, which only moderately affects Crb localization (Das and Knust, 2018) likely due to the continued operation of polarized cytoskeletal transport (Fig 6C-E). Similarly, mutation of the FERM-binding domain of Crb has strong effects upon overexpressed Crb (Fletcher et al., 2012; Letizia et al., 2011) but only causes a moderate reduction of endogenously expressed Crb (Cao et al., 2017; Klose et al., 2013; Sherrard and Fehon, 2015a) (Fig 5E,F). Thus, important mechanisms of Crb polarization that are obscured through genetic redundancy can be revealed through the study of overexpressed Crb to provide a unifying model of polarization (Fig 6F).

Our results explain the striking genetic interactions we have observed between upstream components of the Hippo signaling pathway, whose disruption leads to increased Crb expression (Fletcher et al., 2018; Genevet et al., 2009; Hamaratoglu et al., 2009; Zhu et al., 2015), and components of the Crb trafficking machinery such as exocyst components (Fletcher et al., 2015; Fletcher et al., 2012). Specifically, RNAi knockdown of spectrin cytoskeleton components in *sec15* mutants or *kib* mutants, or alternatively double mutants of *ex kib*, all cause strong Crb accumulation in endosomes in the follicle cell epithelium (Fletcher et al., 2015; Fletcher et al., 2012) (Fig S2). Since Crb itself functions with the spectrin cytoskeleton, the FERM domain proteins Merlin and Expanded, and the exocyst-binding partner Kibra to directly regulate Hippo signaling, follicle cells make use of this pathway to sense mechanical stretching of the apical domain to promote increased expression of *crb, ex, kib* and other target genes to help maintain the apical domain and accommodate mechanical perturbation (Fletcher et al., 2018). Thus, by virtue of being a transmembrane protein, an apical polarity determinant, as well as an upstream component of the Hippo pathway, Crb helps orchestrate cytoskeletal polarization, but is also transported apically, so can act as a sensor of the successful maintenance of cytoskeletal polarity in the face of significant mechanical stress or strain exerted upon the cytoskeleton of epithelial cells during development.

Finally, the model we propose in *Drosophila* epithelia may be conserved in humans, as the subcellular localisation of many of the components we have examined is similar in a simple columnar epithelium of the human gallbladder (Fig S3). In particular, human CRB3 localises to sub-apical tight junctions with PALS1 and PAR6, while MYO5A/B, MERLIN, KIBRA, CAMSAPs, spectrins and exocyst components localise across the entire apical surface (Fig S3). Notably, CRB3 lacks the homophillic extracellular domain of CRB1 and CRB2, but is still able to localise to tight junctions, suggesting that other tight junction proteins such as JAMs, Occludins or Claudins may mediate extracellular domain clustering, with the entire complex clustered by intracellular multi-PDZ domain proteins such as ZO-1 and MUPP1/PATJ (Bazellieres et al., 2018; Ebnet et al., 2004; Michel et al., 2005). In addition, our findings suggest that polarisation of the cytoskeleton in human columnar epithelial cells may also contribute to apically-directed membrane trafficking of CRB1-3 and other transmembrane tight junction components. In future, it will be of interest to test whether the mechanisms of epithelial polarization uncovered in *Drosophila* are conserved in epithelial organoids, an experimentally tractable model system which forms highly columnar cells similar to those observed in human epithelial tissues.

## Materials and Methods

Mitotic clones were generated using the FLP/FRT system and were either marked positively (presence of GFP; MARCM) or negatively (absence of GFP). Third instar larvae were heat-shocked once at 37°C for 1 hour and dissected 3 days after eclosion. Expression of UAS-driven transgenic lines was achieved with *traffic jam.Gal4* (*tj.Gal4)* driver, the actin ‘flip-out’ and MARCM systems. For ‘flip-out’ clones, third instar larvae were heat-shocked at 37 °C for 20 minutes, and dissected 3 days after eclosion. Fly crosses were kept at a temperature of 25 degrees.

### Immunohistochemistry

Ovaries were dissected in PBS, fixed for 20 minutes in 4% paraformaldehyde in PBS, washed for 30 minutes in PBS/0.1% Triton X-100 (PBT) and blocked for 15 minutes in 5% normal goat serum/PBT (PBT/NGS). Primary antibodies were diluted in PBT/NGS and samples were incubated overnight at 4°C. Secondary antibodies were used for 2 hours at room temperature and then mounted on slides in Vectashield (Vector Labs). Images were taken with a Leica SP5 confocal using 40x oil immersion objective and processed with Adobe Photoshop and ImageJ.

Primary antibodies used were: rat anti-Crumbs (1:200, E. Knust), mouse anti-Crumbs (Cq4) (1:10, DSHB), rabbit anti-Lgl (1:50, Santa Cruz), mouse anti-Dlg (1:250, DSHB) and FITC-conjugated anti-GFP (1:400, Abcam).

Secondary antibodies used were goat Alexa fluor 488, 546 or 647 (1:500, Invitrogen), Phalloidin (2.5:250, Life Technologies) to stain F-actin and DAPI (1 μg/ml, Life Technologies) to visualize nuclei.

### Inhibitor treatments

Treatment of ovaries expressing *Crb::GFP* was performed by isolating egg chambers and culturing them as described (Aguilar-Aragon et al., 2018) with Colchicine (0,2 mg/ml), Latrunculin A (0,05 mM), Cytochalasin D (0,05 mM), Jasplakinolide (0,05 mM), Ethanol or DMSO control (all of them from Sigma) for 2 hours. After treatment, samples were fixed and processed normally for imaging.

### Statistical Analysis

Experiments were performed with at least three biological replicates. Prism software was used to plot the mean of the experimental data and error bars represent the standard deviation. T-test for all conditions tested in the paper was found to be p<0.01.

### Drosophila Genotypes

Fig 1A: *w;;crb-GFP*

Fig 1B: *w; tj.Gal4/+; crb-GFP/UAS.dynein-IR*^*(28054 VDRC)*^

Fig 1C: *w hs.flp;; actin.FRT.STOP.FRT.Gal4 UAS.GFP/ UAS.dynein-IR*^*(28054 VDRC)*^

Fig 1D: *w hs.flp;; actin.FRT.STOP.FRT.Gal4 UAS.GFP/ UAS.dynein-IR*^*(28054 VDRC)*^

Fig 2A: *w;;crb-GFP*

Fig 2B: *w; tj.Gal4/+; crb-GFP/UAS.dynein-IR*^*(28054 VDRC)*^

Fig 2C: *w; tj.Gal4/UAS.kinesin-IR; crb-GFP/*

Fig 2D: *w; tj.Gal4/UAS.kinesin-IR; crb-GFP/UAS.dynein-IR*^*(28054 VDRC)*^

Fig 3A: *w; tj.Gal4/+; UAS.MyoV-GFP/+*

> *w; tj.Gal4/+; UAS.GFP-MyoV-GT/+*

Fig 3B: *w*

Fig 3C-D&F: *w; tj.Gal4/+; UAS.GFP-MyoV-GT/+*

Fig 3E&G: *w; tj.Gal4/+; UAS.GFP-MyoV-GT/ UAS.dynein-IR*^*(28054 VDRC)*^

Fig 4A-B: *w*

Fig 4C-D: *w; tj.Gal4/+; UAS.MyoV-GFP/+*

Fig 4E-G: *w;;crb-GFP*

Fig 5A: *w hs.flp FRT19A moe*^*PL106*^*/FRT19A ubi.RFP*

Fig 5B: *w hs.flp FRT19A mer*^*4*^*/FRT19A ubi.RFP; FRT40Aex*^*e1*^*/FRT40A GFP*

Fig 5C: *w hs.flp FRT19A mer*^*4*^ *moe*^*PL106*^*/FRT19A ubi.RFP*

Fig 5D: *w hs.flp FRT19A mer*^*4*^ *moe*^*PL106*^*/FRT19A ubi.RFP; FRT40Aex*^*e1*^*/FRT40A GFP*

Fig 5E: *w hs.flp;; FRT82B crb*^*ΔFBM*^ */ FRT82B ubi.nlsGFP*

Fig 5F: *w hs.flp;; FRT82B crb*^*ΔFBM*^ */ FRT82B ubi.nlsGFP*

Fig 5G: *w hs.flp UAS.GFPnls tub.Gal4/+;;FRT82Bsec15*^*1*^*/FRT82B tub.Gal80*

Fig 5H: *yw hs.flp tub.Gal4 UAS.GFPnls/+; FRT40Asec5*^*e10*^*/FRT40A tub.Gal80*

Fig 6A: *w*

> *w; tj.Gal4/ UAS.patronin-IR*^*(27654 VDRC)*^
>
> *w; tj.Gal4/+; UAS.shot-IR*^*(41858 BLOOMINGTON)*^*/+*
>
> *w; tj.Gal4/+; UAS.myoV-IR/+*
>
> *w; tj.Gal4/+; UAS.GFP-MyoV-GT/+*
>
> *w; tj.Gal4/+; UAS. α-Cat-IR/+*
>
> *w hs.flp UAS.GFPnls tub.Gal4/+;;FRT82B sec15*^*1*^*/FRT82B tub.Gal80*

Fig 6B: *w; tj.Gal4/UAS.Crb-FL*

> *w; tj.Gal4/UAS.Crb-FL UAS.patronin-IR*^*(27654 VDRC)*^
>
> *w; tj.Gal4/UAS.Crb-FL; UAS.shot-IR*^*(41858 BLOOMINGTON)*^*/+*
>
> *w; tj.Gal4/UAS.Crb-FL; UAS.myoV-IR/+*
>
> *w; tj.Gal4/UAS.Crb-FL; UAS.GFP-MyoV-GT/+*
>
> *w; tj.Gal4/UAS.Crb-FL; UAS. α-Cat-IR/+*
>
> *w hs.flp UAS.GFPnls tub.Gal4/+; UAS.Crb-FL/+; FRT82Bsec15*^*1*^*/FRT82B tub.Gal80*
>
> *yw hs.flp UAS.GFPnls tub.Gal4/+; UAS.Crb-FL/+; FRT82Bkib*^*32*^*/FRT82B tub.Gal80*

---

**Figure S1.**
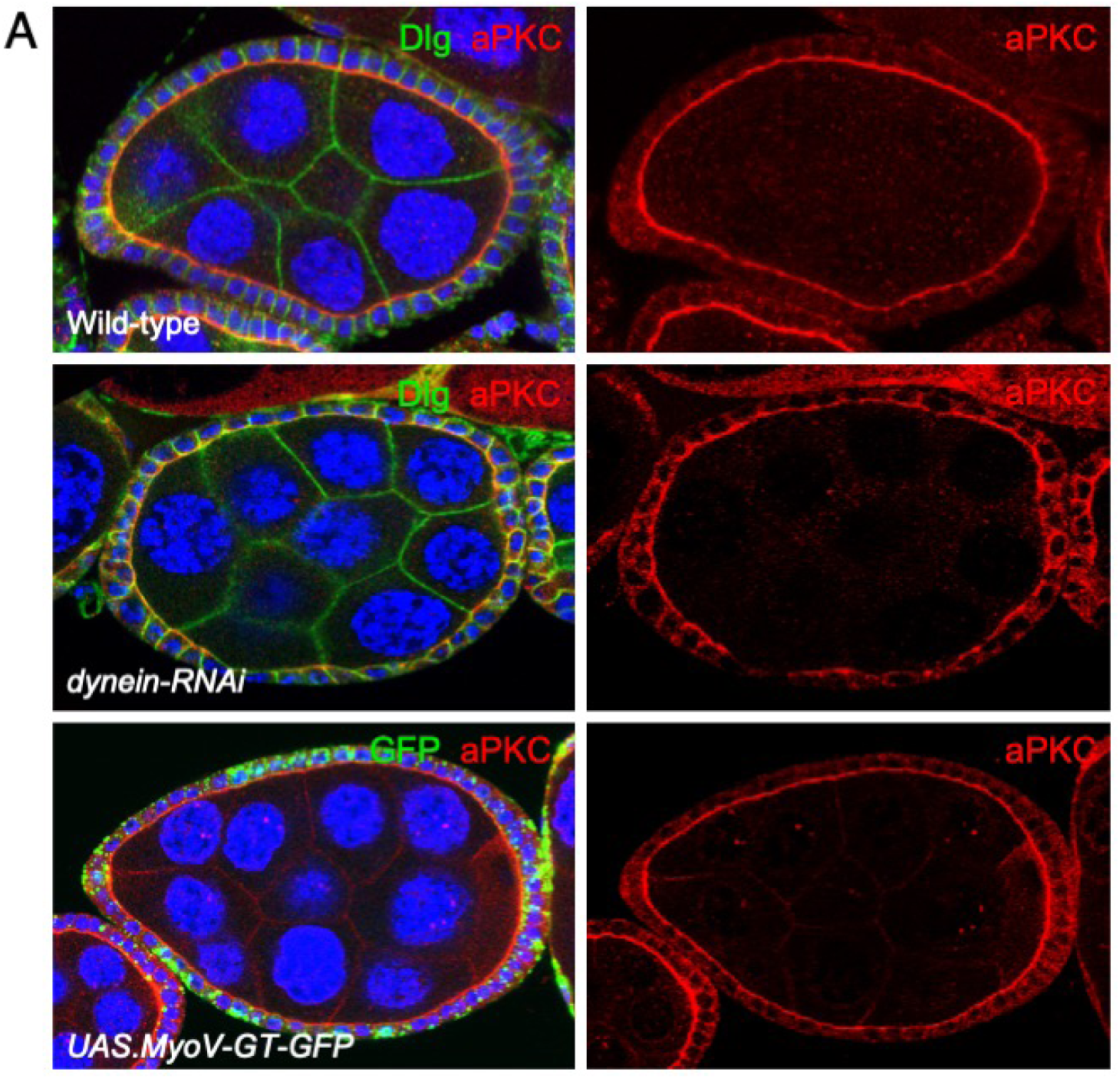
Disruption of apical microtubule or F-actin transport does not affect the polarization of aPKC in the *Drosophila* follicle cell epithelium. A) Egg chambers immunostained for Dig and aPKC show normal apical localization of aPKC even after expression of ***tj.Ga/4 UAS.dynein-RNAi*** or ***tj.Ga/4 UAS.MyoV-GT-GFP.***

**Figure S2.**
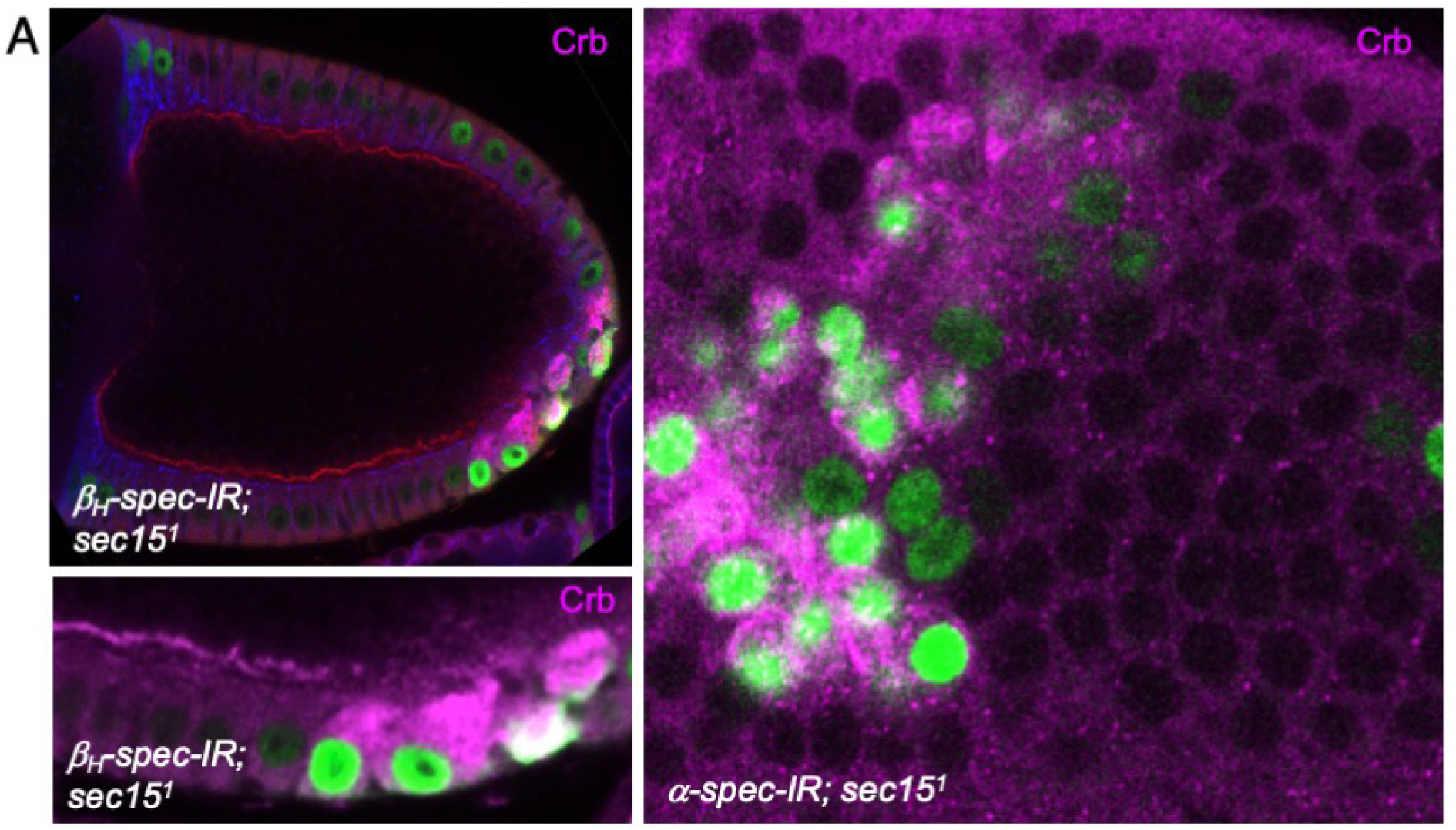
Apical Spectrin acts in parallel with the Exocyst to promote apical delivery of Crb. A) Induction of MARCM clones (GFP+} mutant for *sec15[1]* and expressing *UAS.karst-RNAi* result in massive accumulation of endosomal Crb.

**Figure S3.**
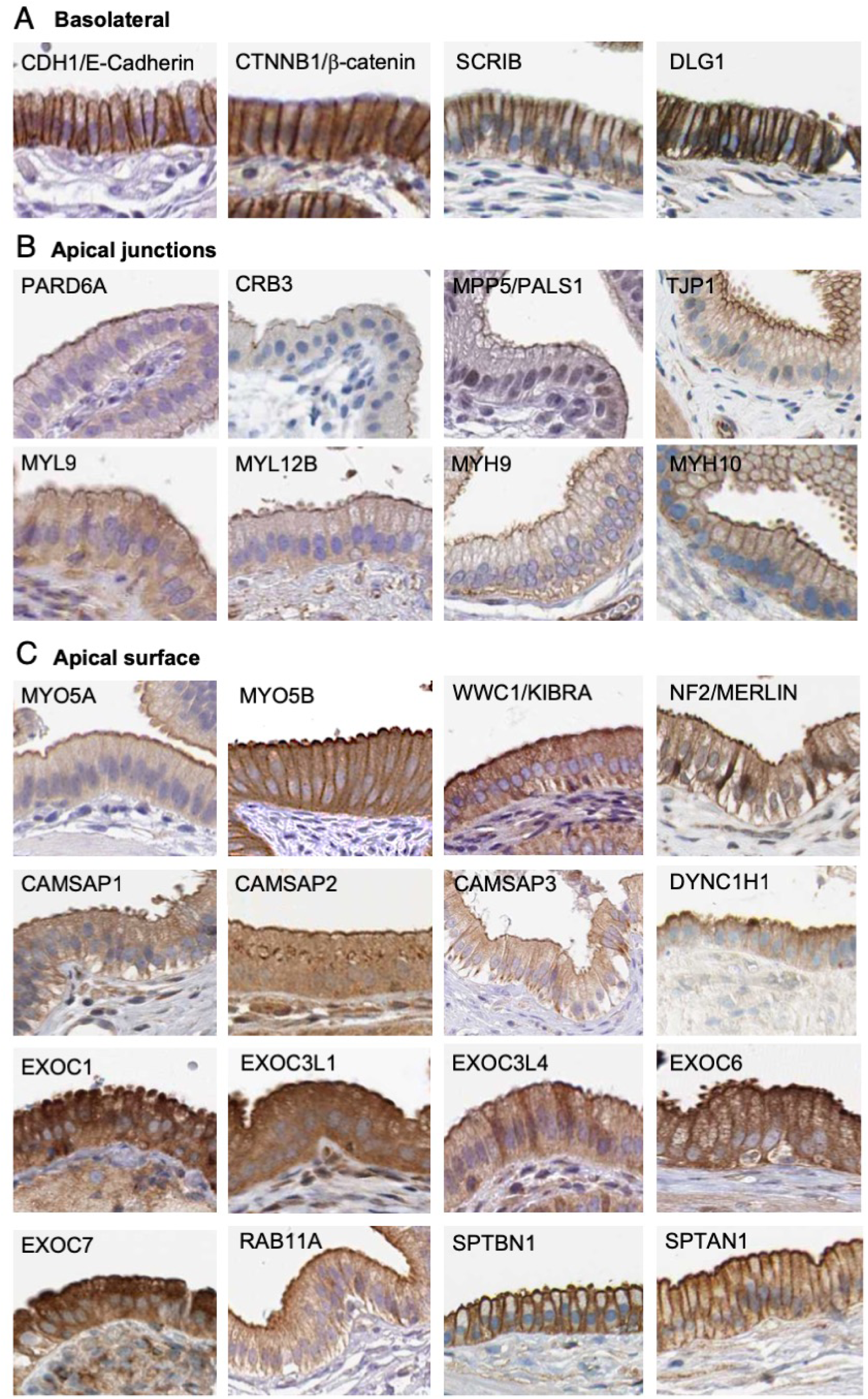
Conserved localization of cytoskeletal polarizing proteins in the human gallbladder epithelium. A) Localisation of basolateral determinants and lateral adherens junction components. B) Localisation of apical proteins that form a ring of apical cell-cell junctions. C) Localisation of apical proteins that spread across the entire apical surface.

